# Fast and reliable quantitative measures of white matter development with magnetic resonance fingerprinting

**DOI:** 10.1101/2024.06.26.600735

**Authors:** Maya Yablonski, Zihan Zhou, Xiaozhi Cao, Sophie Schauman, Congyu Liao, Kawin Setsompop, Jason D. Yeatman

## Abstract

Developmental cognitive neuroscience aims to shed light on evolving relationships between brain structure and cognitive development. To this end, quantitative methods that reliably measure individual differences in brain tissue properties are fundamental. Standard qualitative MRI sequences are influenced by scan parameters and hardware-related biases, and also lack physical units, making the analysis of individual differences problematic. In contrast, quantitative MRI can measure physical properties of the tissue but with the cost of long scan durations and sensitivity to motion. This poses a critical limitation for studying young children. Here, we examine the reliability and validity of an efficient quantitative multiparameter mapping method - Magnetic Resonance Fingerprinting (MRF) - in children scanned longitudinally. We focus on T1 values in white matter, since quantitative T1 values are known to primarily reflect myelin content, a key factor in brain development. Forty-nine children aged 8-13y (mean 10.3y ±1.4) completed two scanning sessions 2-4 months apart. In each session, two 2-minute 3D-MRF scans at 1mm isotropic resolution were collected to evaluate the effect of scan duration on image quality and scan-rescan reliability. A separate calibration scan was used to measure B0 inhomogeneity and correct for bias. We examined the impact of scan time and B0 inhomogeneity correction on scan-rescan reliability of values in white matter, by comparing single 2-min and combined two 2-min scans, with and without B0-correction. Whole-brain voxel-based reliability analysis showed that combining two 2-min MRF scans improved reliability (pearson’s r=0.87) compared with a single 2-min scan (r=0.84), while B0-correction had no effect on reliability in white matter (r=0.86 and 0.83 4-min vs 2-min). Using diffusion tractography, we delineated MRF-derived T1 profiles along major white matter fiber tracts and found similar or higher reliability for T1 from MRF compared to diffusion parameters (based on a 10-minute dMRI scan). Lastly, we found that T1 values in multiple white matter tracts were significantly correlated with age. In sum, MRF-derived T1 values were highly reliable in a longitudinal sample of children and replicated known age effects. Reliability in white matter was improved by longer scan duration but was not affected by B0-correction, making it a quick and straightforward scan to collect. We propose that MRF provides a promising avenue for acquiring quantitative brain metrics in children and patient populations where scan time and motion are of particular concern.

## 1. Introduction

Developmental cognitive neuroscience aims to shed light on evolving relationships between brain structure and cognitive development. To this end, quantitative methods that reliably measure individual differences in the structure of brain tissue are fundamental. Commonly used qualitative MRI sequences, for example, T1-weighted and T2-weighted contrasts, are useful for delineating anatomy and detecting pathologies. However, these qualitative measures do not provide quantitative metrics that reliably index individual differences in tissue properties (Lerch et al., 2017). Qualitative MRI measures are influenced by scan parameters and hardware-related biases (for review, see Cheng et al., 2012) and also lack physical units that can accurately quantify differences among participants. Images also may appear dramatically different due to specific scanner and protocol settings, making it hard to repeat and compare across studies, and within the same individual over time.

In contrast, quantitative MRI (qMRI) techniques aim to quantify specific physical properties of the tissue that shed light on cellular structure. Specifically, quantitative longitudinal relaxation mapping (qT1) has physical units (milliseconds) that are reliable across different hardware setups (Mezer et al., 2013) and have been reported to be primarily sensitive to myelin content in the white matter (Lutti et al., 2014; Stüber et al., 2014; Warntjes et al., 2017). This specificity is crucial to uncover the neurobiological mechanisms of brain development, degeneration, and disease. For example, changes in qT1 values have been described in multiple sclerosis (Harper et al., 2023; Stevenson et al., 2000), reflecting alterations in axon and myelin composition. Similarly, qT1 has been used to study aging (Eminian et al., 2018; Filo et al., 2019), and degenerative and inflammatory processes in clinical conditions (Gozdas et al., 2021; Menke et al., 2009; Moallemian et al., 2023), as well as lifespan maturation and degeneration of white matter (Yeatman et al., 2014).

Although qT1 can provide insights into neurobiological mechanisms, it is not commonly incorporated into studies of child development. Existing qT1 protocols typically require long acquisition times (e.g., Callaghan et al., 2014; Deoni et al., 2005; Gracien et al., 2017; Helms et al., 2008; Kecskemeti et al., 2016; Leutritz et al., 2020; Marques & Gruetter, 2013; Mezer et al., 2013; O’Muircheartaigh et al., 2019; Sanchez Panchuelo et al., 2021), which limits feasibility for children and clinical populations that have difficulties staying in the scanner for long periods of time. Moreover, children tend to move while in the scanner (Barkovich et al., 2019; Greene et al., 2018), which is a major challenge for many quantitative measures that require multiple images to be collected in sequence. For all these reasons, loss of data is a major barrier to large-scale quantitative MRI studies in children. When scans are required for medical reasons, it is a common practice to sedate children in order to get good data quality, though sedation also has health implications (Artunduaga et al., 2021; Jaimes et al., 2018; Wilder et al., 2009) and is therefore rarely used in research. This makes the development of reliable sequences with short scan times of particular importance for pediatric research and practice.

Magnetic resonance fingerprinting (MRF) is an efficient quantitative MRI technique that has the potential to addresses these limitations by providing a rapid and high-resolution data acquisition, particularly with recent advancements that have further improved its acquisition efficiency (Cao et al., 2019, 2022; Liao et al., 2024; Ma et al., 2013, 2018). MRF is designed to simultaneously quantify different tissue properties in a single fast acquisition. With recent improvements it can achieve whole brain 1-mm isotropic mapping in 2 minutes, which paves the way for using quantitative MRI across a broad range of research and, eventually, in clinical practice. While several studies have assessed the reliability and reproducibility of MRF in phantoms and healthy human adults (Buonincontri et al., 2019; Fujita et al., 2023; Gracien et al., 2020; Körzdörfer et al., 2019; Wicaksono et al., 2023), the use of MRF in pediatric studies has been scarce (Chen et al., 2019; Kim et al., 2023; Liao et al., 2024). To date, the reliability and quality of MRF-based measurements in pediatric and clinical populations, where data quality is a major challenge, has never been evaluated.

In the current study, we evaluate the scan-rescan reliability of MRF-estimated T1-maps in a longitudinal pediatric sample. In a group of children scanned several months apart, we show that MRF-derived T1 values are 1) highly reliable, 2) precisely capture individual differences in tissue properties, and 3) replicate known age effects in white matter tracts. In addition, we examine different reconstruction pipelines and evaluate the influence of different pipeline choices on scan-rescan reliability. We conclude with recommendations for incorporating MRF acquisitions in studies of brain development in health and disease.

## 2. Methods

### 2.1. Participants

49 children aged 8-13y (mean 10.3y ± 1.4; 29/20 Female/Male) completed two scanning sessions 2-4 months apart at the Stanford University Center for Neurobiological Imaging, as part of an ongoing longitudinal study. All children provided informed assent to participate in the study and their parents provided written consent. The study protocol was approved by the Stanford University School of Medicine Institutional Review Board. Prior to the first scan, children were acclimated to the MRI environment with a mock MRI scanner. All data were acquired on a 3T GE Discovery MR750 UHP (GE Healthcare, Milwaukee, WI, USA), equipped with a Nova 32-channel head coil.

### 2.2. Data acquisition

#### 2.2.1. MRF acquisition

In each session, we collected two 3D-MRFs and a separate ‘PhysiCal’ calibration scan. MRF data were acquired using tiny golden-angle shuffling MRF with optimized spiral-projection trajectories as proposed in (Cao et al., 2022). In the MRF sequence, an adiabatic inversion-prepared pulse is used with an inversion time of 20 ms, followed by acquisition periods. Each acquisition period contains 500 TRs with varying flip angles, termed an ‘acquisition group’. A resting time of 1.2 s follows each acquisition group to allow for signal recovery before the next acquisition group. Different acquisition groups use different complementary spiral k-space trajectories. A total of 16 acquisition groups are needed in each MRF sequence to obtain 1-mm whole brain quantitative (2 minute scan time). The field of view (FOV) was 220*220*220 mm^3^, with reconstruction matrix of 220*220*200. While the two MRF scans used the same parameters, they included complementary k-space acquisition trajectories to allow them to be combined synergistically to produce a 4-minute scan with reduced k-t undersampling. In addition, a unified, rapid calibration sequence called Physics Calibration (PhysiCal) (Iyer et al., 2020) was used to measure B0 inhomogeneity within the same FOV, with resolution of 2*2*2 mm^3^ in 25 seconds.

#### 2.2.2 Diffusion-weighted acquisition

We collected multi-shell dMRI data with 180 diffusion weighted volumes, distributed across three shells with the following gradient scheme: 30 directions with b=1000 s/mm^2^, 60 directions with b=2000 s/mm^2^ and 90 directions with b=3000 s/mm^2^, as well as 14 reference volumes without diffusion weighting (b = 0 s/mm^2^). We used TR of 3335 ms with minimal echo time to obtain spatial resolution of 1.5 mm^3^ isotropic voxels in 84 axial slices. This was achieved using a hyperband acceleration with a slice acceleration factor of 4. The scan duration totalled 11 minutes and was broken down to two 5.5 minute scans to allow children a short break. An additional scan of six non-diffusion-weighted volumes with a reversed phase encoding direction and the same parameters was also acquired to correct for echo-planar imaging (EPI) distortions.

#### 2.2.3 Anatomical acquisition

A high-resolution T1-weighted (T1w) anatomical scan was acquired using GE’s BRAVO sequence, which is a fast SPGR with a spatial resolution of 0.9mm^3^ isotropic voxels.

### 2.3 Data processing

#### 2.3.1 MRF image reconstruction

##### B0 field inhomogeneity estimation

From the PhysiCal data, highly under-sampled multi-echo images were reconstructed with parallel imaging and compressed sensing methods. Subsequently, a robust multi-echo general linear modeling robustly recovered an artifact-free B0 map.

##### Subspace reconstruction without B0 inhomogeneity correction

The reconstruction used a spatiotemporal subspace modeling with locally low rank (LLR) constraint based on previous literature (Cao et al., 2022). The MRF dictionary was pre-calculated using the extended phase graph method (Weigel, 2015). A singular value decomposition (McGivney et al., 2014) was applied on the pre-calculated dictionary to get the first five temporal principal components, termed as subspace basis Φ. The coefficient maps *c* of the bases can be reconstructed by this formula:

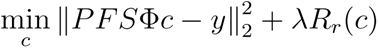

Where *P* is the under-sampling pattern; *F* is the nonuniform Fourier transform (NUFFT); *S* is the coil sensitivity map; *Y* is the acquired k-space data; *R_r_*(*c*) is LLR term and *λ* is the regularization.

##### Subspace reconstruction with B0 inhomogeneity correction

With the presence of B0 field inhomogeneity, the acquired signal becomes *e^itwm^y* and equation (1) becomes

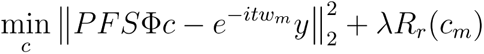

where *e^itwm^y* is the conjugate phase demodulation with acquisition time *t* accumulated during the acquisition trajectory at an off-resonance frequency *w_m_*. In this study, we incorporated a time-segmented B0 correction method (Irarrazabal et al., 1996; Noll et al., 1991) to correct the B0-inhomogeneity induced image blurring.

##### Template matching

Subspace-based compression was also applied to the pre-calculated dictionary, which can significantly reduce the computational load and memory requirements. Then, a cross-correlation template matching was applied to the reconstructed coefficient maps using the compressed dictionary to obtain quantitative T1 values.

##### MRF reconstruction pipelines

In order to assess the scan time effects on image quality, data were reconstructed in four ways: 1) using a single 2-min MRF data. 2) Combining two 2-min MRF scans (4-min). 3) Using a single 2-min MRF in combination with B0 correction. 4) Combining two 2-min MRF scans with B0 correction. Importantly, before reconstructing the 4-min data, we estimated the motion between the two 2-min MRF scans, and corrected the motion in k-space.

#### 2.3.2 dMRI preprocessing

Diffusion data were pre-processed using the default pipeline in QSIprep 0.19.1 (Cieslak et al., 2021). The T1w image was corrected for intensity non-uniformity using *N4BiasFieldCorrection* (Tustison et al., 2010), ANTs 2.4.3) and reoriented into AC-PC alignment. This image was used as an anatomical reference throughout the workflow. The diffusion data were denoised with MP-PCA denoising as implemented in MRtrix3’s *dwidenoise* (Veraart et al., 2016) with a 5-voxel window. After denoising, the mean intensity of the diffusion-weighted series was adjusted so that the mean intensity of the b=0 images matched across each separate DWI scanning sequence. B1 field inhomogeneity was corrected using *dwibiascorrect* from MRtrix3 with the N4 algorithm (Tustison et al., 2010). FSL ’s Eddy (version 6.0.5.1:57b01774) was used for head motion correction and Eddy current correction (Andersson & Sotiropoulos, 2016). Eddy was configured with a q-space smoothing factor of 10, a total of 5 iterations, and 1000 voxels used to estimate hyperparameters. A linear first level model and a linear second level model were used to characterize Eddy current-related spatial distortion. q-space coordinates were forcefully assigned to shells. Eddy’s outlier replacement was run (Andersson et al., 2016). Data were grouped by slice, only including values from slices determined to contain at least 250 intracerebral voxels. Groups deviating by more than 4 standard deviations from the prediction had their data replaced with imputed values. b=0 reference images with reversed phase encoding directions were used along with an equal number of b=0 images extracted from the primary diffusion scans. The susceptibility-induced off-resonance field was estimated using a method similar to that described in (Andersson et al., 2003). The fieldmaps were ultimately incorporated into the Eddy current and head motion correction interpolation. The final interpolation was performed using the jac method. Finally, the two diffusion weighted series were concatenated and resampled to the anatomical AC-PC space with 1.5mm isotropic voxels.

#### 2.3.3. Fiber tractography

Voxel-level diffusion modeling and whole brain tractography were carried out using QSIprep’s MRtrix3 (Tournier et al., 2019) reconstruction pipeline. Within each voxel, diffusion was modeled with constrained spherical deconvolution (CSD) using the *dhollander* algorithm for estimating the fiber response function (Dhollander et al., 2016, 2019). Whole brain tractography was carried out using the iFOD2 algorithm with the multi-shell multi-tissue method (msmt-csd; (Tournier et al., 2004, 2008). One million streamlines were generated and their length was limited to the range 50-250mm, resulting in a whole brain tractogram in each subject’s native space for each timepoint.

pyAFQ (Kruper et al., 2021; Yeatman, Dougherty, Myall, et al., 2012) was then applied to the whole brain tractograms to i) segment white matter tracts of interest and ii) calculate tract profiles along the trajectory of each tract. A standard cleaning procedure was used to remove spurious streamlines that do not follow the trajectory of the desired tract. Specifically, we removed streamlines that deviated from the tract core by more than 3 standard deviations, or that were longer than the mean tract length by more than 4 standard deviations. This process was repeated iteratively 5 times.

Diffusion metrics were then sampled onto 100 equidistant nodes along each tract, such that for each metric the value at the node represents the weighted average across the streamlines that comprise the tract. Here we limit our analyses to the middle 80 nodes of each tract, as nodes closer to the tract terminations typically suffer from partial volume as streamlines enter gray matter. To evaluate reliability across different regions of the brain, we segmented both inter- and intra-hemispheric tracts. First, we segmented the 7 sub-bundles of the corpus callosum (CC): Orbital, Superior Frontal, Motor, Superior Parietal, Posterior Parietal, Temporal and Occipital (Dougherty et al., 2007). Then, we also segmented 7 intra-hemispheric tracts that have been frequently studied in developmental cognitive neuroscience: the direct and posterior branches of arcuate fasciculus (ARC, pARC), the superior longitudinal fasciculus (SLF), the inferior fronto-occipital fasciculus (IFOF), the inferior longitudinal fasciculus (ILF), the uncinate fasciculus (UF), and the corticospinal tract (CST). Each tract was segmented bilaterally (see Figure 1).

**Figure 1.**
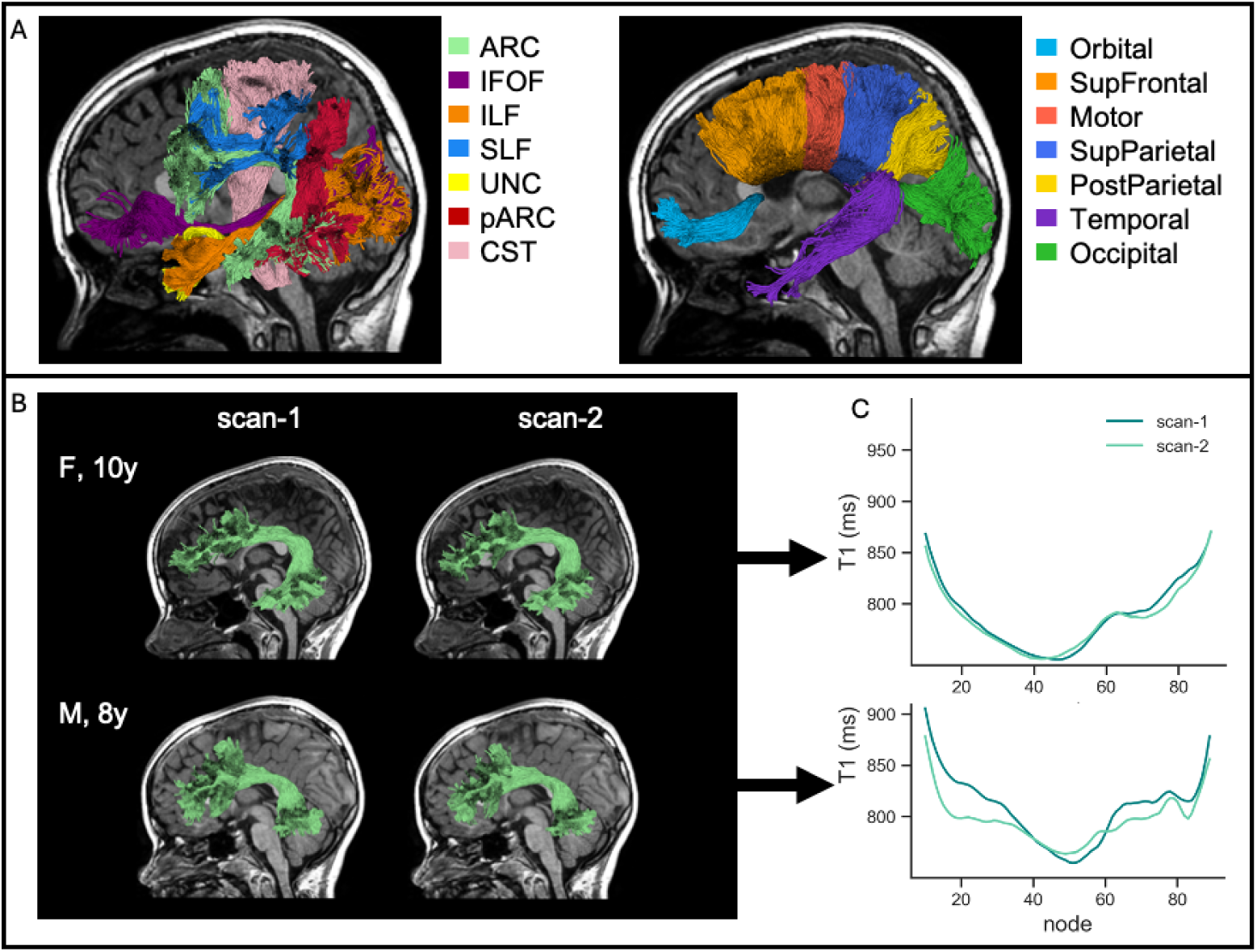
Calculating T1 profiles along the trajectory of major white matter tracts. A) Intra-hemispheric tracts (left) and callosal sub-bundles (right) of a representative subject, overlaid on their anatomical T1w image. B) Left Arcuate Fasciculus (ARC) of two example subjects across two timepoints. Tract shape is variable between subjects yet highly consistent within the same subject over time. C) T1 profiles along the trajectory of the left ARC.

### 2.4 Reliability analysis

We took three approaches to evaluate scan-rescan reliability in white matter: (1) voxel-wise analysis across the entire white matter, (2) ROI-based analysis of the corpus callosum (CC, and (3) tractometry-based comparisons in multiple white matter tracts (Figure 1). In each of these three analyses we compared values obtained from the first scan available for each child to the values obtained in their second scan, which took place 2-4 months apart. This provides a lower limit on scan-rescan reliability as there might be some developmental changes over this time-scale.

#### 2.4.1 Voxel-wise and ROI-based comparison

Accurate registration of the two sessions is an essential step for assessing scan-rescan reliability. We took several steps to achieve high quality registration (see Figure 2): First, we registered and resampled the two scans into a ‘half-way’ space, to maintain inverse consistency instead of resampling the source (the second scan) to the estimated target location (the first scan) (Reuter et al., 2010; Reuter & Fischl, 2011). Second, instead of registering MRF-generated T1 maps, we registered the reconstructed coefficient maps before template matching, which reduces the partial volume effects and thus improves the accuracy of T1 fitting (Chen et al., 2023). The registration was calculated for the 4-min reconstruction pipeline and then applied to all pipelines. Then, to obtain accurate white matter masks in the ‘half-way’ space, the two ‘half-way’ space 4-min coefficient maps were averaged to generate a higher SNR image (equivalent to an 8-min scan), followed by template matching and synthetic T1w-MPRAGE generation. This high quality image was used as input for the Freesurfer segmentation pipeline using the SynthSeg method (Billot et al., 2023), to obtain the white matter mask and corpus callosum (CC) ROIs (see Figure 2). CC ROIs were visually inspected and manually edited to exclude corticospinal fluid (CSF) and fornix voxels.

**Figure 2:**
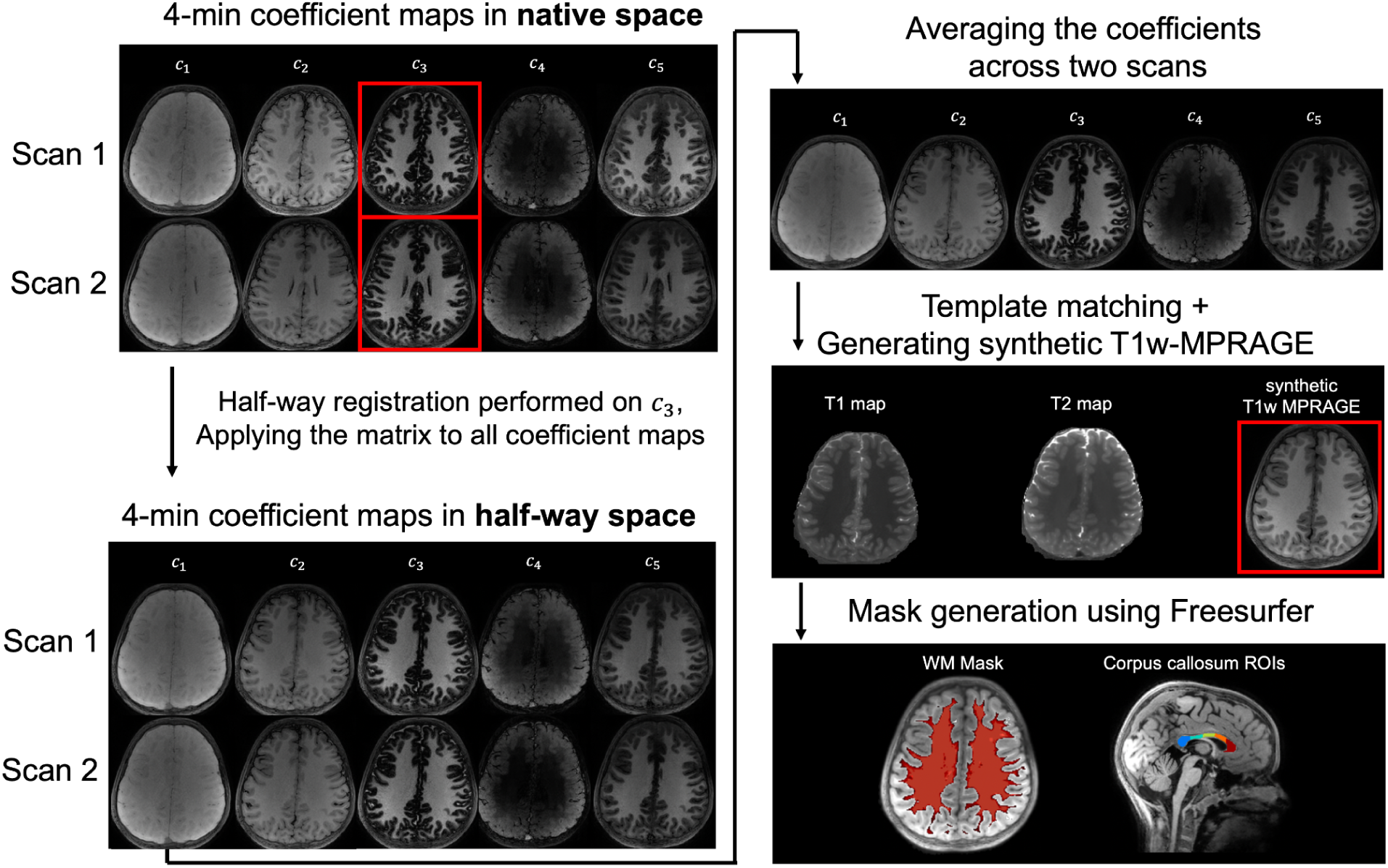
Workflow for the half-way registration process. Initially, 4 minute coefficient maps from two scans are presented in native space, where the *c_3_* coefficient (highlighted in a red box) was chosen as the registration input due to its high contrast. Then, we applied the calculated transform matrices from each scan to the half-way space to all five coefficient maps. We then averaged the registered coefficient maps from both scans to increase SNR, before running the template matching to generate synthetic T1-weighted MPRAGE images. Lastly, masks were created using Freesurfer software for white matter and corpus callosum regions of interest.

The scan-rescan reliability was first assessed voxel-wise within the entire white matter mask. Pearson’s correlation coefficient was calculated among voxels of the two scan sessions for each participant. Coefficient of determination (R^2^) was further calculated with the fitting function y=x to assess the data consistency. We ran a linear mixed effect model using the lme4 package in R (Bates et al., 2015) to evaluate the contribution of scan duration and B0-correction to scan-rescan reliability. The model included a random intercept for each subject to account for individual variability.

For the ROI-based analysis, we focused on five corpus callosum regions as the CC is an area with dense, highly myelinated axons (Aboitiz et al., 1992). The average T1 value in each ROI was calculated per subject and compared across the two scanning sessions. Again, we used Pearson’s correlation and coefficient of determination (R^2^) across all subjects as measures of scan-rescan reliability.

#### 2.4.2 Tractography based comparison

In order to evaluate T1 values along the tracts, we first had to register the MRF data to the native diffusion space. To achieve this, synthetic diffusion b=0 images were generated based on the T1 and T2 maps from the 4-min MRF scan. These synthetic images were registered to the diffusion b=0 images using Advanced Normalization Tools (ANTs) (Avants et al., 2009; Klein et al., 2009). A rigid linear transformation was chosen with optimizing a mutual information (MI) similarity metric. The transformation was subsequently applied to the MRF coefficient maps, followed by template matching to get T1 values mapped into the diffusion native space. T1 values obtained from each of the four different pipelines were then sampled onto 100 equidistant nodes along each tract using pyAFQ.

We evaluated scan-rescan reliability by comparing the mean T1 values of each tract across the two scans of each participant. We calculated Pearson’s correlation coefficient and coefficient of determination (R^2^) between the two scans, when T1 values were estimated using each of the four pipelines.

### 2.5. Age effects

To examine whether MRF-derived values are able to replicate known developmental effects (Yeatman et al., 2014), we calculate Pearson’s correlation between mean R1 values in each tract and participants’ age at the time of each scan. For this analysis we used R1, the inverse of T1 (1/T1), as our metric, since R1 linearizes the T1 scale and has been more extensively studied in relation to brain development (Travis et al., 2019; Yeatman et al., 2014). This was repeated for the four pipelines to test whether different processing choices affect the ability to detect expected age effects.

## 3. Results

### 3.1 Qualitative evaluation of reconstructed maps

Figure 3 shows two typical slices of reconstructed qT1 maps from the four different pipelines. All reconstructions resulted in 1mm resolution images with good contrast between gray matter, white matter and CSF. Compared with the 2-min acquisition, the 4-min acquisition resulted in less noisy maps, irrespective of B0-correction (first row of Figure 3A). B0-correction significantly reduced image blurring around the sinuses, which are air-filled spaces in the skull that induce high B0-inhomogeneity (Figure 3A).

**Figure 3.**
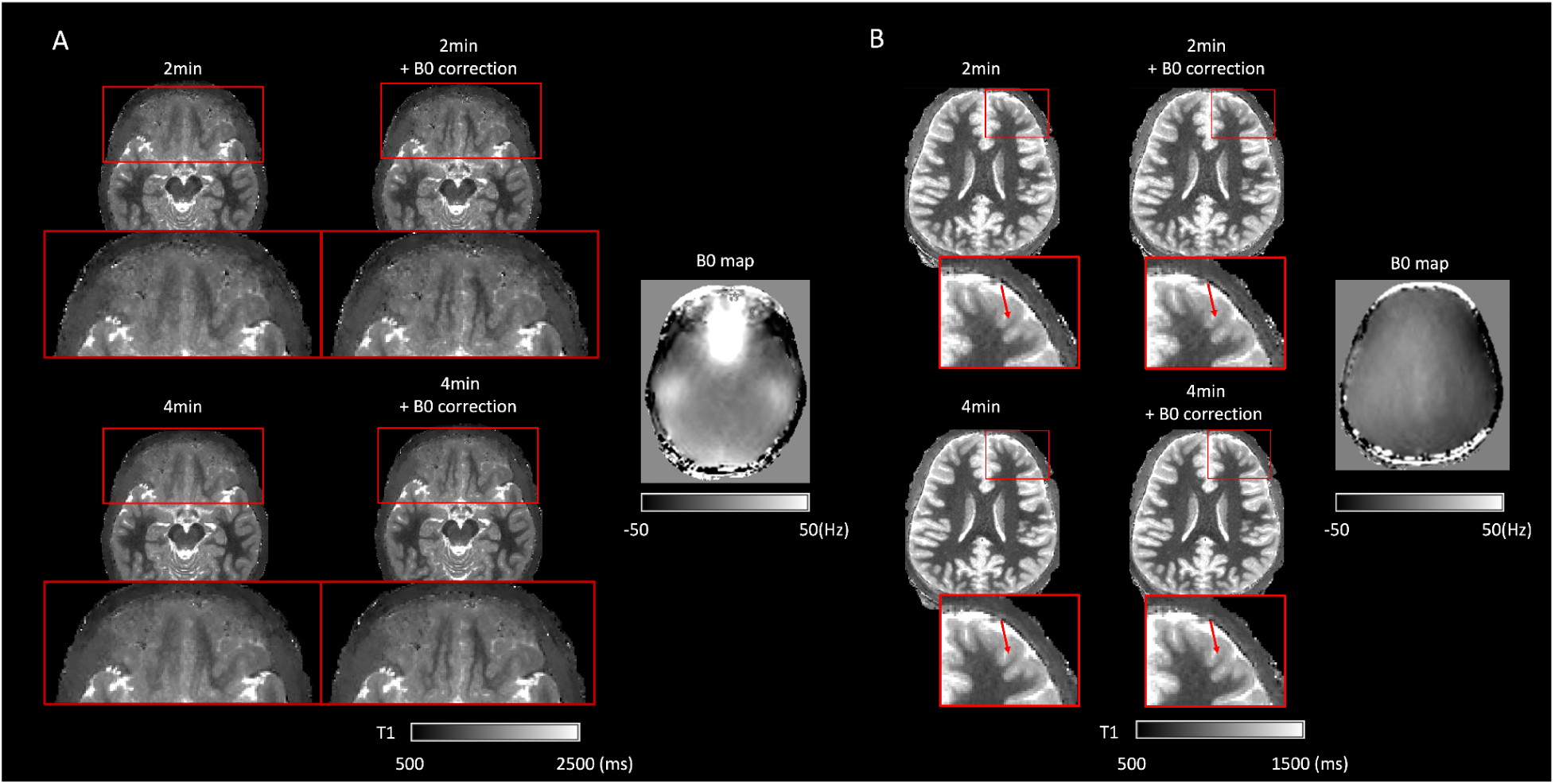
Example axial slices of reconstructed T1 maps using the four pipelines. A) A slice passing through the sinuses, a region with high B0 inhomogeneity (as shown in the corresponding B0 map on the right). Applying B0 correction significantly sharpens the image in the inhomogeneous area. B) A more superior slice where B0 is more homogenous. Here, B0 correction does not show significant improvement to image quality in white matter. In both slices, maps generated from 4-min data are less noisy than those of 2-min data.

### 3.2 Voxel-wise comparison

Figure 4 shows scan-rescan reliability across all white matter voxels in a single subject (panel A) and the distribution of correlation coefficients across all participants in the sample (panel B; one subject was removed from the analysis due to severe blurring caused by motion). Individual participants showed high Pearson’s correlation for the 2-minute data (r = 0.838 ± 0.041, range [0.744 - 0.918]) and coefficient of determination (R^2^ = 0.638 ± 0.114, range [0.278 - 0.835]), indicating that individual variation in qT1 values across voxels is highly reliable across scans. As expected, reliability was higher for 4-minute data, suggesting that longer scan duration improved SNR (r = 0.866 ± 0.043, range [0.772 - 0.927]; R^2^ = 0.693 ± 0.121, range [0.319 - 0.853]). B0-correction did not improve reliability compared to data without this correction (2-minute: r = 0.838 vs r = 0.834, R^2^= 0.638 vs 0.629 for without vs with B0-correction; 4-minute: r = 0.866 vs 0.863, R^2^: 0.693 vs 0.683 for without vs with B0-correction). A linear mixed effect model confirmed that reliability was higher for 4-min data compared with the 2-min data (β = 0.028, t = 12.85, p < 1e-16), but was not affected by the B0-correction (β = -0.004, t = -1.705, p = 0.09). The interaction between duration and B0-correction was not significant (p = 0.95). Lastly, to examine reliability in the absence of developmental changes, we also compared the values of the two 2-minute scans that were collected at the same scanning session. As expected, scan-rescan reliability was higher within session compared with the 2-minute data across sessions (r = 0.841 ± 0.031, range [0.696 - 0.880]; R^2^ = 0.673 ± 0.065, range [0.365 - 0.755]).

**Figure 4.**
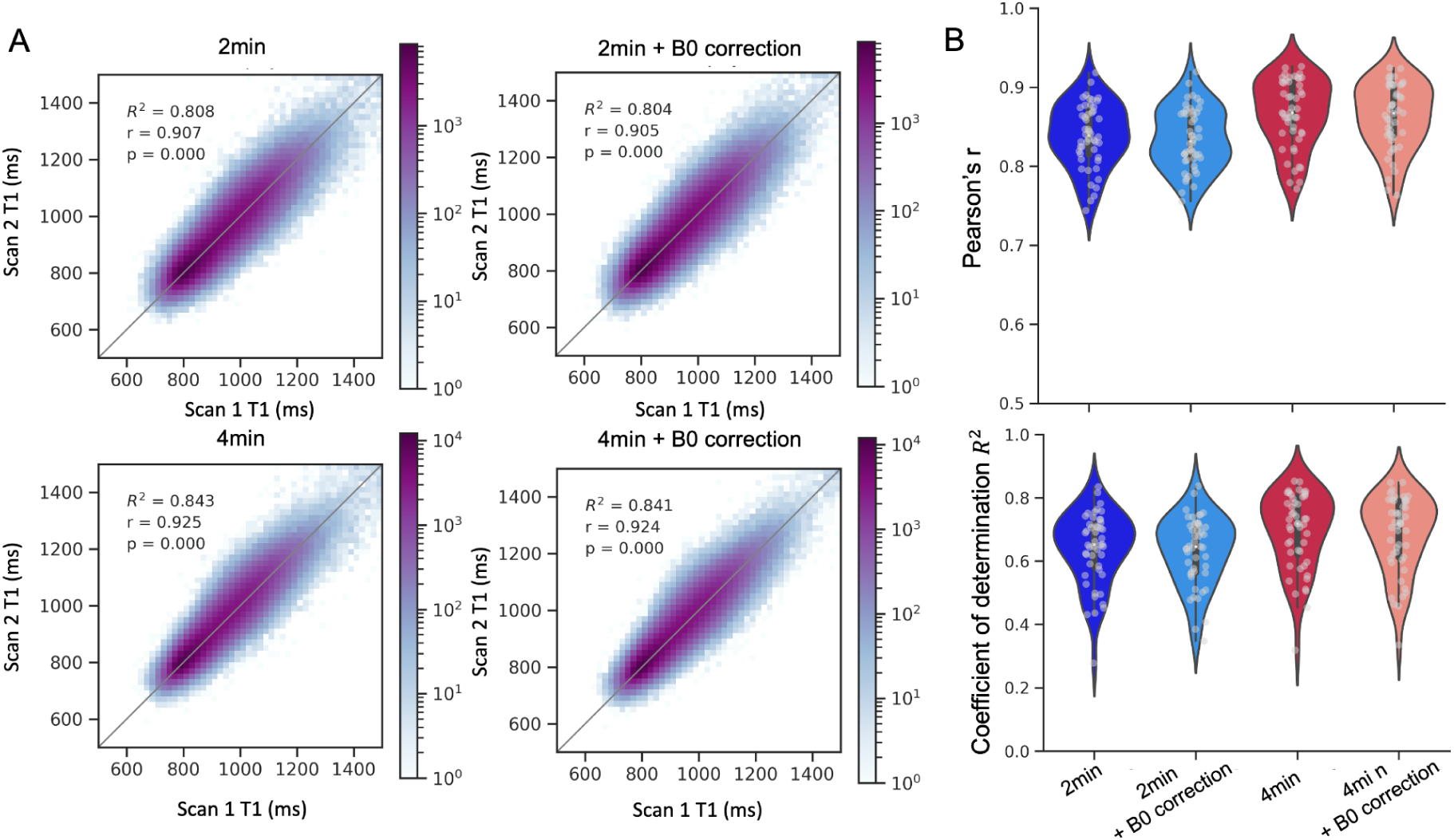
Voxel-wise scan-rescan reliability. A) 2D Histograms show T1 values in white matter voxels in the two scans of a single subject, for the four reconstruction pipelines. Color denotes voxel count in logarithmic scale. Gray lines denote the equality line. B) scan-rescan reliability across all subjects (N=48), calculated as pearson’s correlation (top panel) or coefficient of determination (bottom panel).

### 3.3 ROI-based comparison

We next evaluated mean T1 values in five segments of the corpus callosum (CC), obtained in each subject’s halfway space using Freesurfer. We first observed that T1 values follow an inverted U-shape (Figure 5B), in line with previous findings using traditional qT1 methods and histology (Aboitiz et al., 1992; Berman et al., 2018; Hofer et al., 2015; Lynn et al., 2021; Stikov et al., 2011, 2015). Across the 5 segments, T1 values were highly reliable for all four reconstruction pipelines, with slightly higher reliability for 4-min data (Figure 5C). Similarly to the voxel-based analysis, B0-correction did not contribute to reliability in the CC segments.

**Figure 5.**
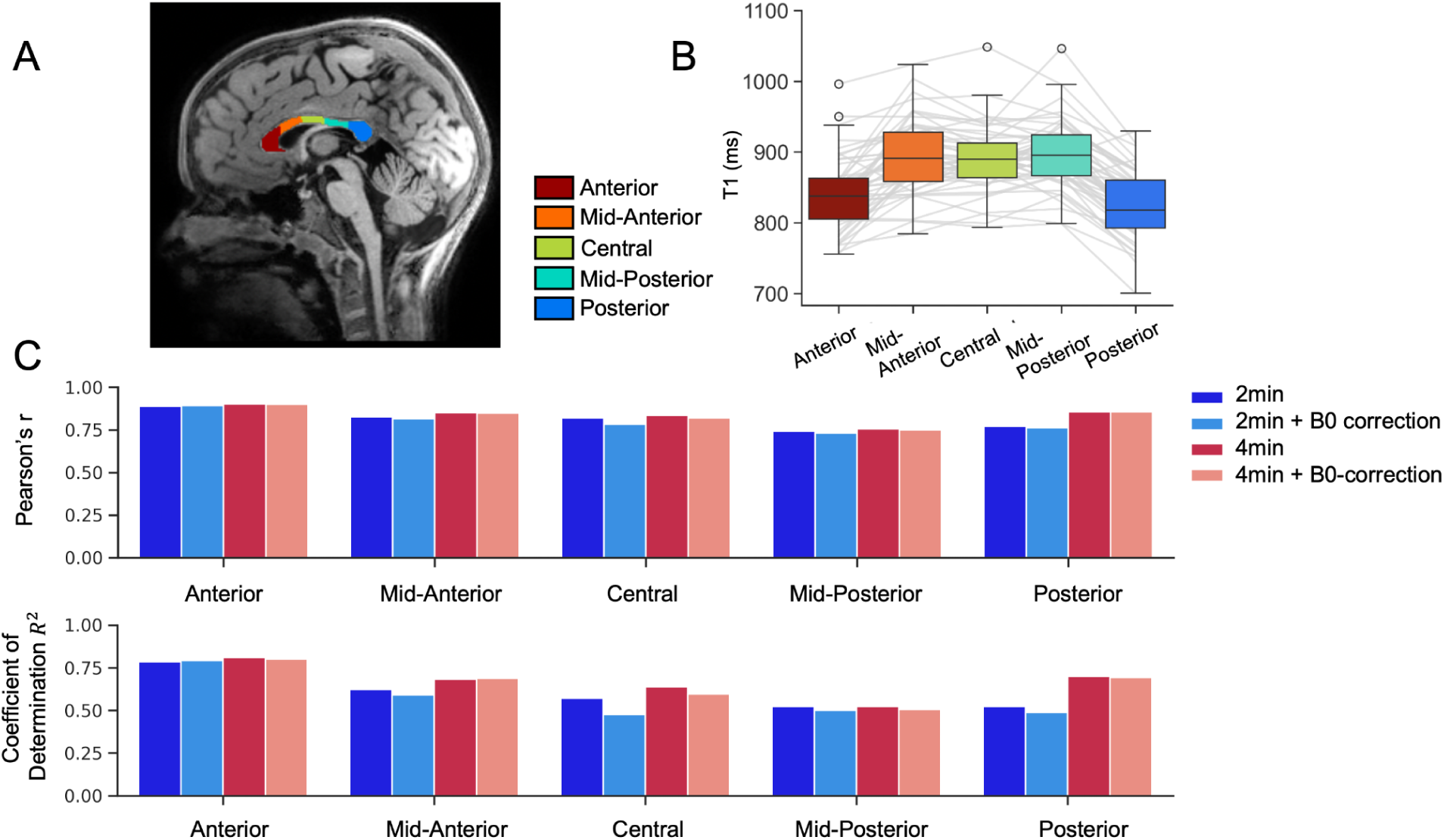
scan-rescan reliability and T1 values in the Corpus Callosum (CC) A). Five regions of interest (ROIs) of the CC overlaid on a synthetic T1w image of a representative subject. B) T1 values exhibit an inverted U-shape along the anterior-posterior axis of the CC. Gray lines denote individual subjects. C) scan-rescan reliability in the CC ROIs is high across all pipelines, with subtle improvement for the 4-min pipelines.

### 3.4 Reliability of T1 values in white matter tracts

We excluded high-motion diffusion scans where the mean framewise displacement (FD) was greater than 0.8mm, or the neighborhood correlation was lower than r = 0.6. This resulted in a sample size of 41 subjects that had good quality diffusion data in both timepoints.

We first evaluated scan-rescan reliability by comparing mean tract T1 values across the two scans of each participant, calculated using each of the four different pipelines. This analysis showed that across all tracts, the reliability was high even for the single two minute scan without B0 correction (mean Pearson’s r = 0.827 across 21 tracts, range [0.751-0.887]). Increasing the scan duration to 4 minute slightly increased reliability (mean r = 0.836, range [0.709-0.898]), while B0 correction hardly had an effect on reliability (mean r = 0.826, range [0.753-0.887]; mean r = 0.836, range [0.712-0.897] for two and four minutes, respectively). These results are consistent with what we reported above for the voxel-based and ROI-based analyses. Figures 6, 7, and Supplementary Figure S1 show the variability in scan-rescan correlation values across different tracts, and Supplementary Table 1 reports the corresponding coefficients of determination.

**Figure 6.**
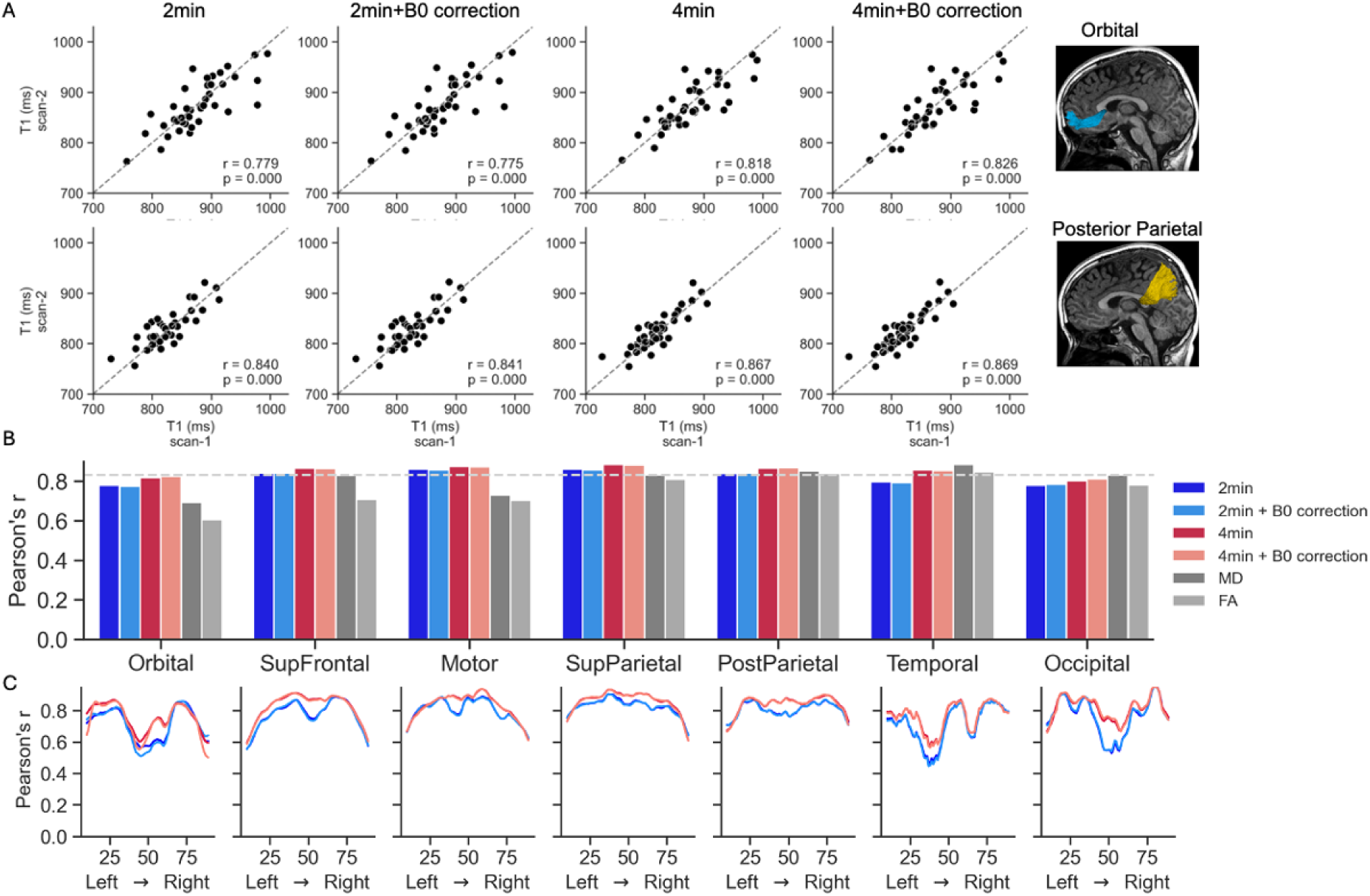
scan-rescan reliability of callosal white matter tracts using the 4 reconstruction pipelines. A) Mean T1 values for the first and second scan of each participant in the orbital sub-bundle (top) and posterior parietal sub-bundle of the corpus callosum (CC). Dashed lines represent the equality line. B) Pearson’s r correlation coefficient for mean T1 values across the 4 pipelines for the seven CC sub-bundles. Diffusion metrics are shown for reference in gray (MD, mean diffusivity; FA, fractional anisotropy). Dashed line represents the median reliability across all tracts. C) Reliability along the tract profile. In each sub-bundles, nodes are ordered from left to right.

For reference, we also calculated the reliability of diffusion-based metrics for each tract, fractional anisotropy (FA) and mean diffusivity (MD), in the same manner. This analysis revealed that MRF-derived T1 values show comparable or higher reliability to that of widely used diffusion-based metrics (FA: mean r = 0.754 range [0.584 - 0.904; MD: median r = 0.795 range [0.548 - 0.926]). As shown in Figures 6-7 (and supplementary Figure S1), the improved reliability of MRF compared to diffusion stands out especially for tracts where diffusion metrics have relatively low reliability, like the Uncinate Fasciculus and Corticospinal tract.

**Figure 7.**
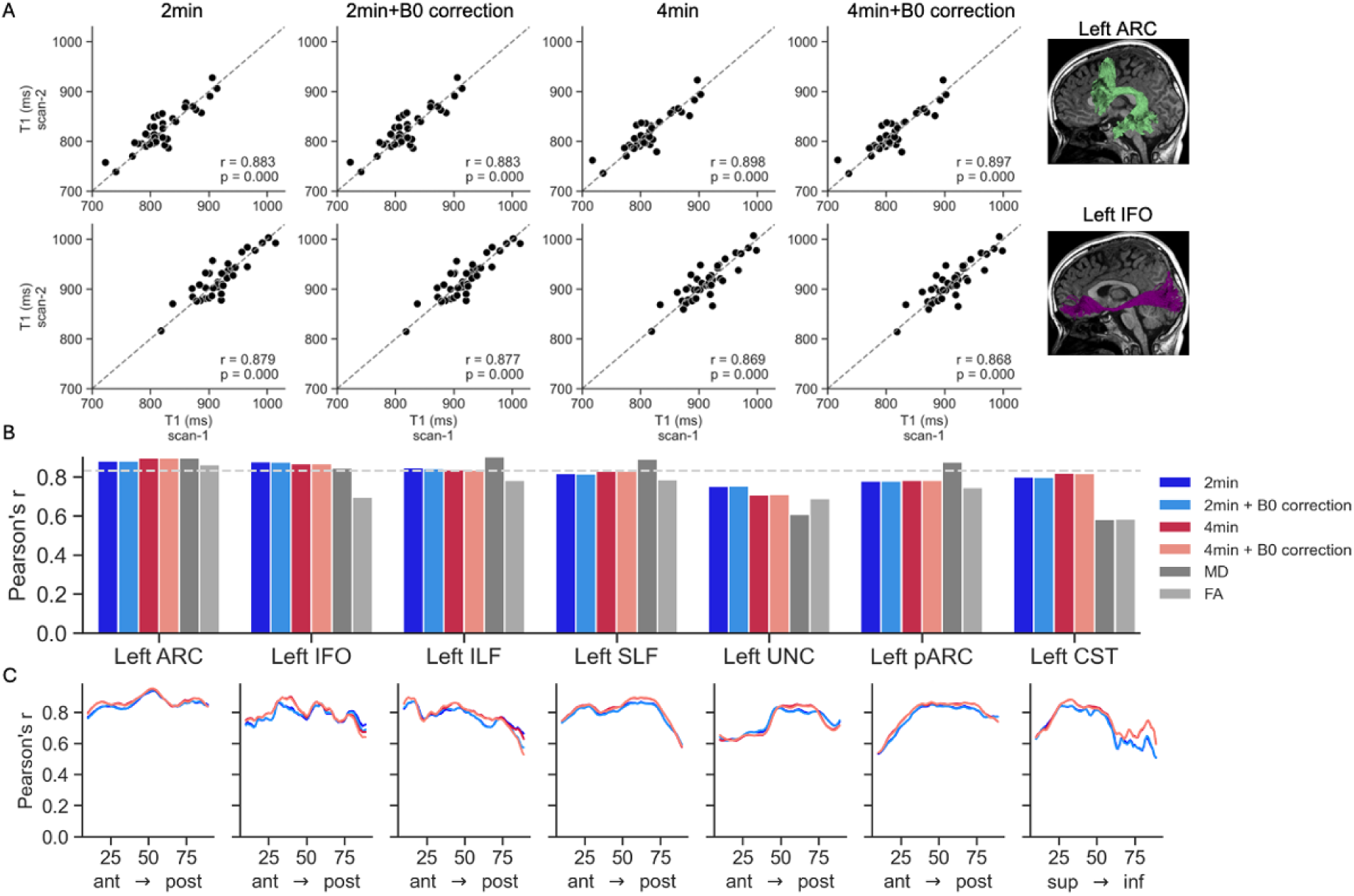
scan-rescan reliability of left hemisphere white matter tracts using the 4 reconstruction pipelines. A) mean T1 values for the first and second scan of each participant in the left arcuate fasciculus (ARC, top) and left inferior fronto-occipital fasciculus (IFO, bottom). Dashed lines represent the equality line. B) Pearson’s r correlation coefficient for mean T1 values across the 4 pipelines in left hemisphere tracts. Diffusion metrics are shown for reference in gray (MD, mean diffusivity; FA, fractional anisotropy). Dashed line represents the median reliability across all tracts. C) Reliability along the tract profile. In each tract, nodes are ordered from anterior to posterior position.

Lastly, we examined whether R1 (1/T1) values along different tracts replicate known age effects. We found that R1 was positively correlated with age in multiple intra-hemispheric tracts, as well as in the more frontal sub-bundles of the corpus callosum (Figure 8 and supplementary Figure S2), in line with previous findings (Yeatman et al., 2014). Repeating this analysis using the four pipelines revealed similar results. Interestingly, there was a trend where in some tracts age effects were larger when R1 values were calculated with 4-minute pipelines, compared with the 2-minute pipelines. While not statistically significant, this might suggest that using 4-min sequences increases SNR and enhances the ability to capture underlying effects of tissue development.

**Figure 8.**
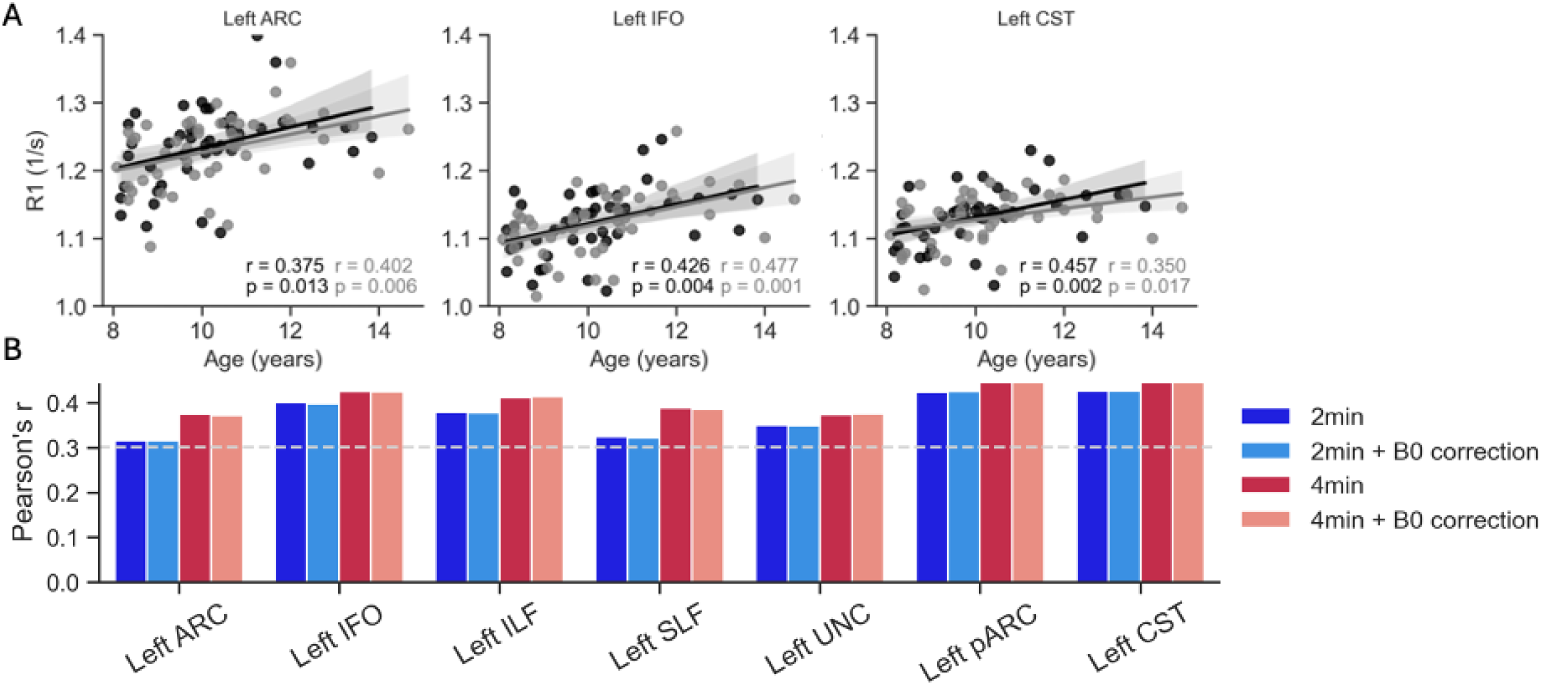
Tract mean R1 values positively correlate with age. A) Scatterplots show R1 values calculated using the 4-minute pipeline (without B0 correction) from the two scans of each participant (first scan N=43, black; second scan N=46, gray). Lines denote the best linear fit surrounded by the 95% confidence interval (shaded area). Shown are the left Arcuate Fasciculus (ARC), left Inferior Fronto-Occipital Fasciculus (IFO) and left corticospinal tract (CST). B) Correlation coefficients between age and R1 values in left hemisphere tracts when calculated using the four different pipelines (scatterplots in A correspond to the red bars). The dashed line denotes uncorrected p<0.05 as a reference.

## 4. Discussion

In this study, we evaluated the reliability and validity of a novel and rapid MRF sequence that can be used to obtain high quality qT1 data in young children. Our analyses showed that MRF-derived T1 values were highly reliable in white matter over time, and were able to capture known age effects. In addition, we evaluated the contribution of different acquisition and preprocessing choices on scan-rescan reliability.

Our analyses revealed that high scan-rescan reliability in white matter can be achieved with a single two minute scan and that reliability increases with scan. While previous studies evaluated the reliability of MRF in phantoms and healthy adults (Buonincontri et al., 2019; Gracien et al., 2020; Körzdörfer et al., 2019; Wicaksono et al., 2023), here we conducted a more realistic assessment of reliability in a real life research scenario with young children. Our results provide a proof of concept for the feasibility of using MRF to measure tissue properties reliably over time with populations where data acquisition and quality are more challenging. These results hold important implications for the field of developmental cognitive neuroscience and pave the way for studies that focus on structural brain development in children.

We examined the reliability of MRF-derived T1 values in white matter using three complementary methodologies, namely, comparing individual voxels, analyzing specific ROIs, and using diffusion based tractography. This three-pronged approach was intended to test MRF-derived values in the common analyses used to study white matter structure. Specifically, tractography has been extensively used to study the relationship between white matter development and cognitive functions (Roy et al., 2024; Wang et al., 2017; Yeatman, Dougherty, Ben-Shachar, et al., 2012). We show that using a standard automated pipeline for tractography, reliability of MRF-derived tract profiles is remarkably high across the tracts examined, and in some cases exceeded the reliability of diffusion-based metrics.

In addition to establishing the reliability of the MRF sequence, the current work validates MRF-derived T1 values in two ways. We first found that T1 values in the corpus callosum follow an inverted U-shape, in line with prior observations using other quantitative methods and histology (Aboitiz et al., 1992; Berman et al., 2018; Hofer et al., 2015; Lynn et al., 2021; Stikov et al., 2011, 2015). This demonstrates that MRF-derived T1 values are sensitive enough to capture this variation in adjacent parts of the corpus callosum. In addition, we observed significant correlations between age and R1 values in multiple intra-hemispheric white matter tracts, replicating known age effects (Callaghan et al., 2014; Eminian et al., 2018; Yeatman et al., 2014). Interestingly, the only callosal sub-bundles that correlated with age were the anterior ones, in line with (Yeatman et al., 2014). Since R1 has been shown to be sensitive to myelin content (Lutti et al., 2014; Stüber et al., 2014), the observed increase in R1 through development is thought to reflect the prolonged myelination of white matter which lasts into adulthood. In our age range, we were able to capture linear growth in R1 values, which suggest that white matter tracts are more heavily myelinated in older children compared with younger children. Together, this shows that MRF-derived T1 values capture similar effects to standard T1 values acquired with longer acquisitions.

One of the principal strengths of this work lies in its implementation within a longitudinal pediatric study, which offers promise for quantitative MRI adoption in brain development research. A central challenge in imaging young children is motion during scans, which significantly compromises data quality and leads to high rates of data exclusion. The proposed MRF protocol overcomes this barrier by employing a fast and robust sequence, which is less likely to be affected by motion. Using short sequences not only mitigates motion artifacts in acquired data, but also increases the probability of acquiring the data successfully, as it is not uncommon for children to end scanning sessions prematurely before completing the full sequence. With a protocol that takes 2-4 minutes, this risk is greatly reduced compared with standard protocols that take about 20 minutes to complete and require a sequence of images. This may also reduce the drop-out rate in follow-up visits which is a major challenge in longitudinal studies. Based on the pipeline comparison we conducted, below we provide a list of practical recommendations for researchers and clinicians interested in using MRF sequences in research and practice.

### 4.1. Practical recommendations

#### 4.1.1. Scan duration

Our findings show that reliability increased when we combined two 2-minute sequences compared with a single run. Importantly, even for the single 2 minute sequence, scan-rescan reliability was greater than r = 0.7, and for white matter tracts was comparable or better than the scan-rescan reliability of diffusion metrics. We recommend that if time allows, two repetitions will boost SNR and reliability, but for clinical studies and situations where time is a limiting factor, a single two minute sequence could provide usable qT1 data for many applications.

#### 4.1.2. B0 correction

Our findings do not indicate a significant improvement in reliability of white matter T1 values by including a separate B0-correction. B0 correction will likely be more important for studies that target orbital regions in the vicinity of the sinuses, which may suffer more from inhomogeneity (see Figure 2). B0-correction may also prove crucial for accurate T2 maps, but this remains to be assessed in future work.

#### 4.1.3. Registration

In order to accurately evaluate the reliability of MRF-derived qT1 we created a half-way space template image and simultaneously resampled both scan sessions into the half-way space (Reuter et al., 2010; Reuter & Fischl, 2011). The common practice of registering one scan to another creates an inherent asymmetry in the sense that the second scan is aligned, registered and resampled to match the spatial configuration of the first scan, while the initial scans remain unaltered. The disparity introduces a systematic bias into the repeatability analysis, such that the values may differ depending on which scan serves as the reference image and which scan is undergoing the registration and resampling. We propose that registration to a half-way space resolves this asymmetry and leads to more accurate results.

### 4.2. Limitations and future directions

The current protocol and preprocessing pipeline do not provide information about motion during the scan. The evaluation of motion was done qualitatively based on the reconstructed images. Quantifying motion would allow us to explicitly quantify the effect of motion on MRF-derived T1 values, and potentially explain some of the variance in scan-rescan reliability across subjects. Another challenge for implementing this protocol widely in clinical settings is that the reconstruction phase is currently lengthy and requires heavy computations. Future work can improve the efficiency of the reconstruction making it feasible in clinical settings (Zhou et al., 2024) while also including online motion correction (Nurdinova et al., 2024).

## 5. Conclusion

In sum, MRF provides a promising methodology for deriving reliable quantitative metrics of brain tissue structure in children and patient populations where scan time and motion are of particular concern. Coupled with the pipeline we propose which improves the repeatability, the current work paves the way for using MRF in longitudinal pediatric studies. This method would facilitate studies focusing on structural brain development by providing an easy and rapid way to quantify changes in myelin and other properties of white matter.

## Acknowledgements

This work was supported by NICHD R01-HD095861 to JDY, grant R01-MH116173 to KS and by the Stanford GSE/SOM postdoctoral fellowship to ZZ. We thank Adam Kerr and Hua Wu from the Stanford CNI for their support and advice in implementing the protocol. We thank Jamie Mitchell, Hannah Stone, Mia Fuentes-Jimenez and Jasmine Tran for their invaluable part in data collection.

## Notes

### Competing Interest Statement

The authors have declared no competing interest.

