## Supplementary Figures for "Fast and reliable quantitative measures of white matter development with magnetic resonance fingerprinting"

### Supplementary material

| | Coefficient of Determination ($R^2$) | | | |
|---|---|---|---|---|
| **Tract** | **2min** | **2min + B0** | **4min** | **4min + B0** |
| ARC_L | 0.765 | 0.765 | 0.799 | 0.797 |
| IFO_L | 0.757 | 0.755 | 0.738 | 0.738 |
| ILF_L | 0.673 | 0.665 | 0.657 | 0.651 |
| SLF_L | 0.623 | 0.619 | 0.664 | 0.664 |
| UNC_L | 0.529 | 0.531 | 0.462 | 0.465 |
| pARC_L | 0.553 | 0.550 | 0.569 | 0.570 |
| CST_L | 0.609 | 0.603 | 0.655 | 0.650 |
| ARC_R | 0.766 | 0.766 | 0.781 | 0.780 |
| IFO_R | 0.736 | 0.740 | 0.690 | 0.692 |
| ILF_R | 0.737 | 0.736 | 0.676 | 0.674 |
| SLF_R | 0.683 | 0.684 | 0.664 | 0.666 |
| UNC_R | 0.491 | 0.490 | 0.521 | 0.529 |
| pARC_R | 0.680 | 0.681 | 0.644 | 0.641 |
| CST_R | 0.603 | 0.600 | 0.670 | 0.669 |
| Orbital | 0.573 | 0.565 | 0.659 | 0.663 |
| SupFrontal | 0.692 | 0.691 | 0.744 | 0.740 |
| Motor | 0.731 | 0.727 | 0.756 | 0.752 |
| SupParietal | 0.719 | 0.713 | 0.767 | 0.765 |
| PostParietal | 0.674 | 0.675 | 0.721 | 0.723 |
| Temporal | 0.625 | 0.619 | 0.733 | 0.729 |
| Occipital | 0.498 | 0.512 | 0.587 | 0.606 |

**Supplementary Table S1.** Coefficient of determination comparing mean T1 values in 21 white matter tracts, across two timepoints using each of the four pipelines.

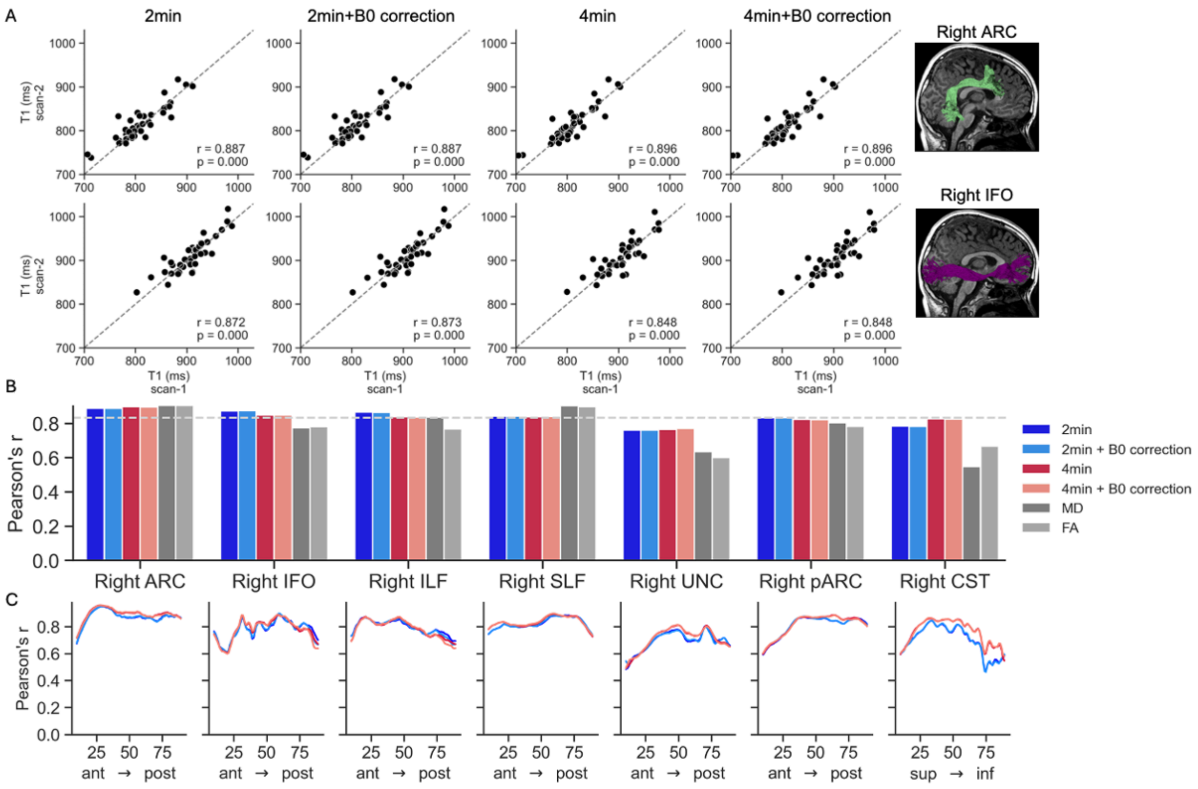


**Figure S1. scan-rescan reliability of right hemisphere white matter tracts using the 4 reconstruction pipelines.** A) mean T1 values for the first and second scan of each participant in the right arcuate fasciculus (ARC, top) and right inferior fronto-occipital fasciculus (IFO, bottom). Dashed lines represent the equality line. B) Pearson's R correlation coefficient for mean T1 values across the 4 pipelines in right hemisphere tracts. Diffusion metrics are shown for reference in gray (MD, mean diffusivity; FA, fractional anisotropy). Dashed line represents the median reliability across all tracts. C) Reliability along the tract profile. In each tract, nodes are ordered from anterior to posterior position.

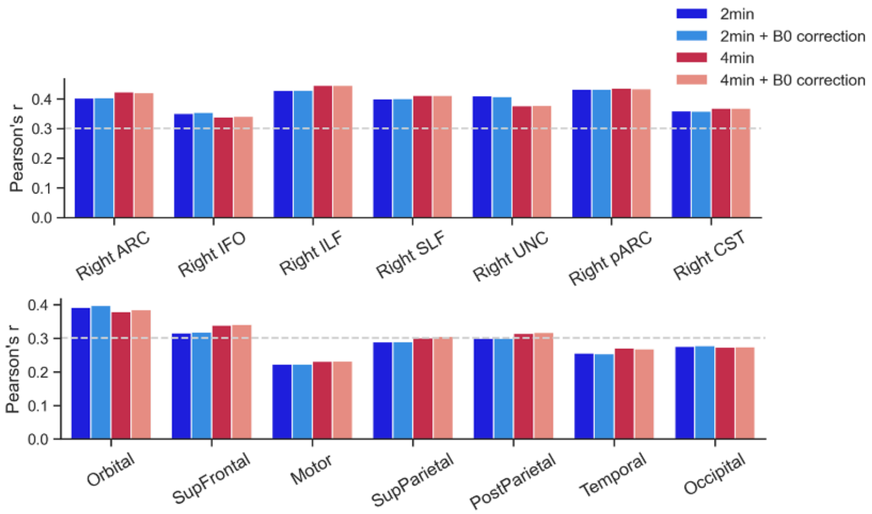


**Figure S2. Correlation coefficients between age and mean R1 values calculated using the four different pipelines, in right hemisphere tracts (A) and callosal sub-bundles (B).** This figure parallels Figure 8B in the main text. The dashed line denotes uncorrected $p<0.05$ for reference.
